# Peptidoglycan recognition protein S2 is crucial for activation the Toll pathway against Israeli acute paralysis virus infection in honey bee *Apis mellifera*

**DOI:** 10.1101/2022.03.02.482613

**Authors:** Yanchun Deng, Sa Yang, Hongxia Zhao, Ji Luo, Zhiqiang Lu, Chunsheng Hou

## Abstract

Although honey bee responses to pathogens have been systematically described in the past decades, antiviral signalling pathways mechanisms are not thoroughly characterized. To decipher direct antiviral roles of an immune pathway, we firstly used the infectious clone of Israeli acute paralysis virus (IAPV) to screen 42 immune genes involved in mTOR, MAPK, Toll, Endocytosis, Jak-STAT pathway and homeobox protein, heat shock protein, as well as antimicrobial peptides (AMPs), and found that Toll pathway was a potential predominant immune pathway in *Apis mellifera*. Consistent with this, only dsRNA-PGRP-S2 treated *A. mellifera* significantly exhibited impaired activation of Toll pathway, promoting susceptibility to the IAPV infection. Finally, immunofluorescence results confirmed that the Toll pathway was initiated by peptidoglycan recognition protein S2 (PGRP-S2) interacting with Toll protein. Co-immunoprecipitation findings also further preliminarily confirmed PGRP-S2 directly interacting with viral capsid protein IAPV-VP3 to induce the activation of the Toll pathway in *A. mellifera*. These findings highlight that the Toll pathway is demanded efficient inhibitions of IAPV replication as a specific antiviral pathway in *A. mellifera*, and PGRP-S2, acting as a pattern recognition receptor, could be a new approach for control of the viral disease.

**Author summary:** Honey bee viruses, particularly IAPV, had been implicated in the colony decline with a global distribution resulting in insufficient pollination services. However, little is known about the antiviral mechanism of honey bee. In this study, we found that the Toll pathway was required for *A. mellifera* against IAPV infection and initiated by PGRP-S2. We also confirmed that dsRNA-PGRP-S2 treated *A. mellifera* exhibited impaired Toll pathway activation and promoted susceptibility to the IAPV infection. As a result, we employed co-immunoprecipitation technique to identify the interaction between the PGRP-S2 with Toll. Moreover, it was found the PGRP-S2 directly recognized IAPV-VP3 to activate the immune pathway against IAPV infection. Our work provides novel evidence that honey bees own a specific antiviral immune pathway and suggests that targeting PGRP-S2 could be a new approach for controlling the viral disease.

## Introduction

Insects rely on a series of immune mechanisms to effectively defend themselves against multiple invading microorganisms. These mechanisms include melanization pathway, autophagy, RNA interference (RNAi), Jak-STAT (the Janus kinase-signal transducer and activator of transcription), Imd (Immune deficiency) and Toll pathway [1]. All of the above pathways are activated via the interaction between the pattern recognition receptors (PRRs) and the pathogen-associated molecular patterns (PAMPs) that are unique to a specific class of microorganisms, and then induce a serial response to eliminate and/or hamper the infection. One of the important PRRs is the Toll-like receptor, which was first identified in *Drosophila*; and is now considered a universal pathogen recognition system [2].

Toll pathway is a well-documented immune response in *Drosophila*, which encoded multiple Toll receptors [3]. Bringing PAMP to Toll results in cleavage of the Toll ligand, spätzle, and then degraded cactus, leading to the expression of antimicrobial peptides (AMPs) [4]. Initially, Toll and Imd pathways are responsible for identifying and clearing Gram-positive/negative and fungi [5]. Recently, Toll pathway in *Drosophila* was shown to be important in innate immune defence against *Drosophila* X virus infection [6, 7]. Subsequently, the Toll pathway was verified to resist the oral viral infection in *Drosophila* [8]. In other insects, such as *Bombix mori*, it had been reported the Toll pathway plays a conserved function as antiviral immune response [9], but little is known about its exact functions in the honey bee antiviral immune. Even though one Toll-like receptor of honey bee, *Am18w*, has been identified, its functions associated with immune response have not been ultimately confirmed yet [10].

Compared with *Drosophila*, studies are lagged on the molecular basis of activation on immune pathways of honey bee. Viruses have caused many losses in bee colonies worldwide [11]. One of the major contributors to great loss was a pathogenic infection caused by RNA viruses [12]. Like other social insects, honey bees have evolved multiple defence mechanisms that counteract the damage induced by RNA viruses at the individual or colony level [13]. Broad antiviral responses depend on RNA interference (RNAi), which was considered as a conserved antiviral defense mechanism in plants and insects [14, 15]. Canonical immune pathways including JAK-STAT, Toll and Imd confirmed their functions against viruses [16]. Chen et al. identified the most up-regulated genes related to immune pathways of honey bee after Israeli acute paralysis virus (IAPV) infection, these pathways include JAK-STAT and Mtogen-activated protein kinase (MAPK) pathway [17]. In addition, the siRNA pathway response to an IAPV infection in *Bombus terrestris* has been described, but they found that siRNA was insufficient to combat viral infection [18]. Transcriptomic and epigenomic analyses have suggested that IAPV infection induced alternation of genes related to immune and epigenetic pathways, these immune genes including *hymenoptaecin*, *PGRP-S2*, *PGRP-S1*, *NF-kappa-B inhibitor cactus 1*, *serine protease 14*, *Toll* and *Raf kinase.* etc. [19]. Recent studies have shown that IAPV caused the changes in most immune genes and pathways (mainly including Toll/Imd pathway and apoptosis) overlapping with those of fruit flies [20]. Despite extensive study on infection characterization of honey bee virus, activation of immune pathways has yet to be fully defined, and how honey bee recognized the viral infections remained unclear. Therefore, the identification of a specific immune pathway to viral infection in honey bee would be the critical step to answer the key questions: (1) is there any immune pathway that is specific for antiviral infection in honey bee?; and if yes, (2) how is it activated and which genes are involved in it?

Most honey bee viruses are positive single-strand RNA viruses in the family *Dicistroviridae* including several important bee viruses such as IAPV. IAPV had been implicated in the colony decline with a global distribution, resulting in insufficient pollination services [17]. Especially, IAPV was found that it had a higher prevalence in colonies failing to overwinter and had been considered a sign of colony collapse disorder (CCD) [17]. In addition, IAPV is one of the best-characterized small RNA viruses and has served as a representative virus in studying the pathogenesis of viral infection in honey bee.

In this study, we used IAPV infectious clone constructed by our laboratory to screen the immune genes involved in the significant immune pathways of honey bee on a large scale [21]. Our results showed that Toll pathway was mainly activated for defending against viral infection of honey bees. A short-type peptidoglycan recognition protein (PGRP-S2) was essential for viral induction of the Toll pathway and activation of AMP genes of *defensin 1* and *hymenoptaecin* to combat the IAPV infection. Our results also revealed that PGRP-S2 was an essential pattern-recognition protein required upstream of the Toll pathway, which was a route-specific to resist viral infection in honey bees. Overall, these findings demonstrated a specific and novel mechanism by which the honey bee elicits immune responses against IAPV infection through activating the PGPR-S2/Toll/AMPs signalling pathway.

## Results

### IAPV Infection Predominantly Induced Expressions of the Toll Pathway associated genes

The previous study has shown that honey bees harboured their antiviral mechanisms [22]. Therefore, the IAPV infectious clone was used to avoid the potential interference from other viruses and explore the precise role in response to viral infection [23]. Our results showed that IAPV infectious clone could effectively reduce honey bee survivals and achieve IAPV proliferation (**S1 Fig**). Then, according to the reported immune genes and pathway against IAPV in Honey bee (**Table S1**) [17, 19, 24], we investigated the expression levels of representative genes involved in immune pathways, including mTOR, MAPK, Toll, endocytosis, Jak-STAT pathways and homeobox protein, heat shock protein genes, as well as AMPs of the honey bee. Analysis of immune genes by qRT-PCR showed that 6 out of 14 genes were significantly up-regulated and the rest were down-regulated in all treated groups (**Fig 1**). Our results showed that IAPV infection elevated ≤ 2-fold the expression of *stat* of JAK-STAT pathway but induced the down-regulation in gene of mTOR pathway (**Fig 1**), which were consistent with that of bees naturally-IAPV-infected [17]. Among these 6 obviously up-regulated genes, *peptidoglycan recognition protein S2* (PGRP-S2), *homeobox protein Hox-C6b*, *defensin 1*, *lysozyme 1* and *hymenoptaecin* were up-regulated at least more than two times, even higher. Especially, *PGRP-S2*, *hymenoptaecin* and *defensin 1* related to Toll pathway were enhanced more than 4-fold after the IAPV treatment. Furthermore, *PGRP-S2* was significantly increased over the survey period in all groups treated with IAPV (*p* < 0.01). In contrast, *apisimin* and *defensin 2* were down-regulated more than 8 times.

**Fig 1.**
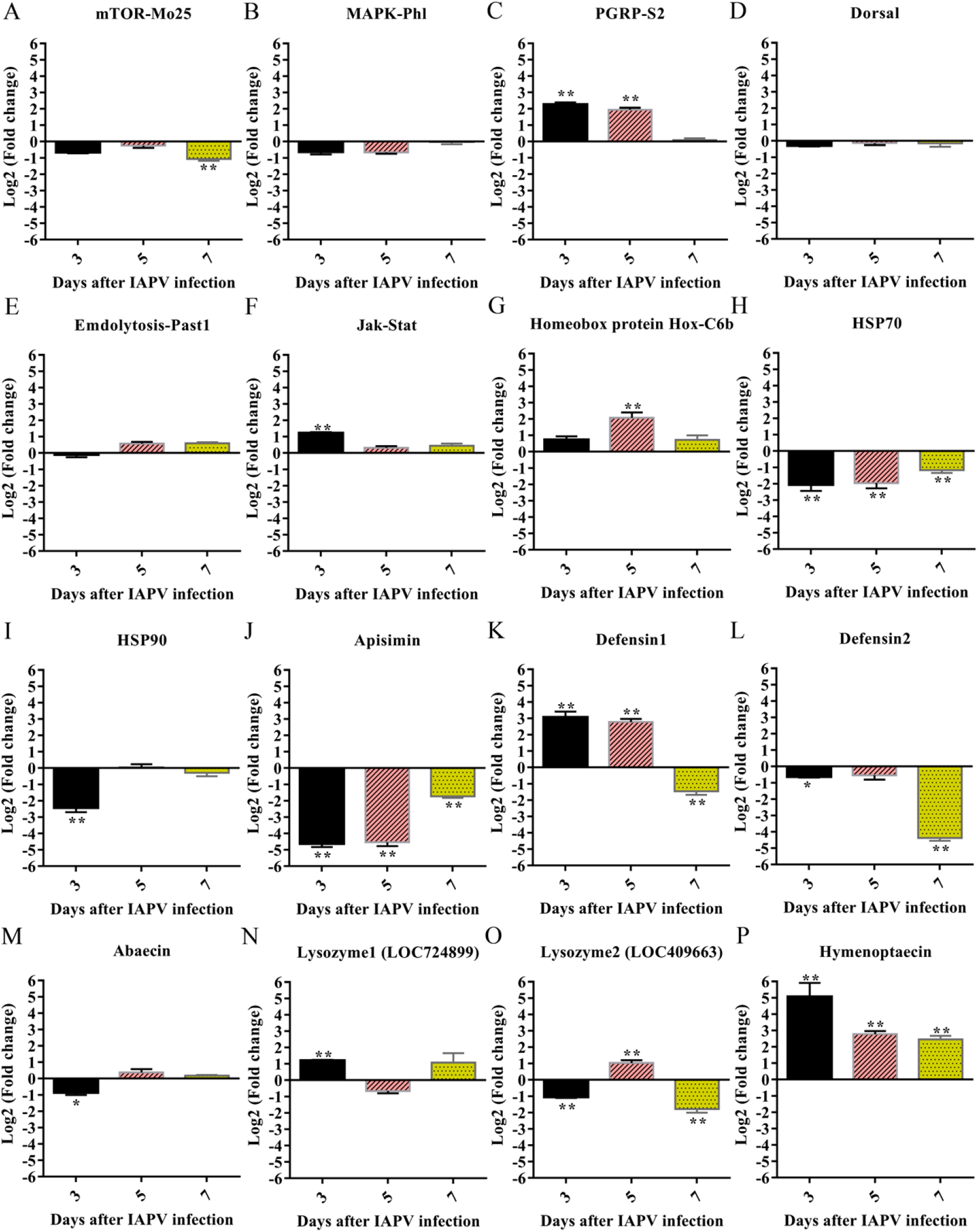
IAPV Infection Elicits the Change of Expression of Immune Genes in Bees. The fold changes in gene expression for mTOR (A), MAPK (B), Toll (C, D), Endocytosis (E), Jak-STAT (F) and homeobox protein, heat shock protein genes (G, H, I), as well as AMPs (J, K, L, M, N, N, O, P). Asterisks indicate significant differences relative to controls (\**P* < 0.05, \*\**P* < 0.01).

The results obtained above raised the issue of the immune signalling pathway involved in regulating the response to viral infection. Then, we focus on investigating the expression levels of the 20 genes involved in the Jak-STAT, RNAi, melanization pathway, especially the Toll and Imd pathway. As showed in **Fig 2 A–D**, most genes of the Toll pathway including *PGRP-S2*, *Toll*, *Dorsal* and *cactus 1* were significantly upregulated after IAPV infection (*P<0.05*), yet *PGRP-LC* and *relish* genes from Imd immune pathways, as well as other genes from Jak-STAT, RNAi, melanization pathway, and two heat shock proteins, had no significant difference before and after viral infection (**Fig 2E–T**). A comparison with a previously published list of 274 overlap immune genes against IAPV infection from Galbraith *et al*. [19] and Li-Byarlay *et al*. [20] was represented, and the only two the significant pathways, Toll signalling pathway (*P <0.00001*) and MAPK signalling pathway (*P <0.01*) were significantly enriched (**Fig 3A**). These results showed that the Toll signalling pathway might be the IAPV-specific inducible immune response.

**Fig 2.**
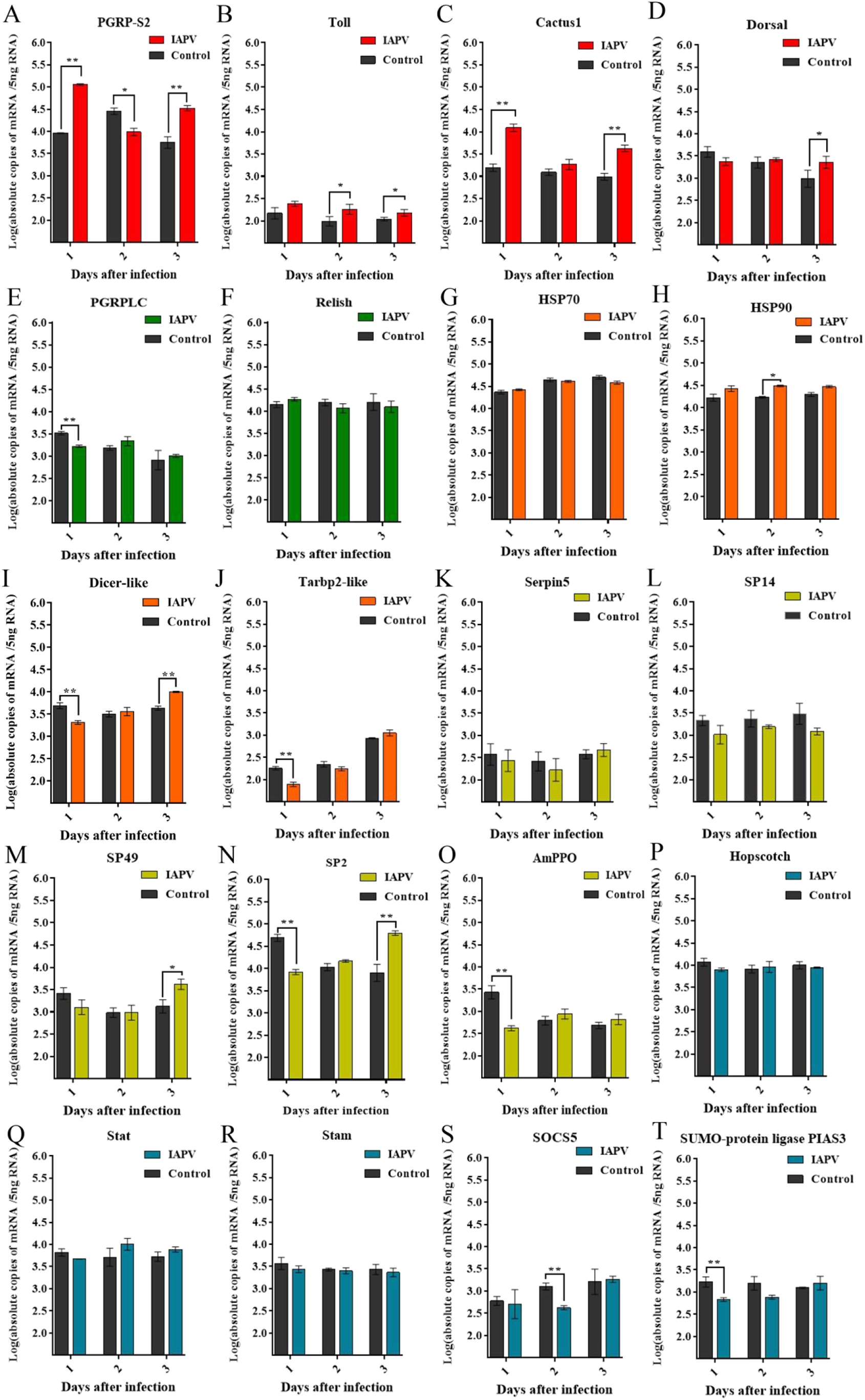
The Toll Pathway was Responsible for Fighting IAPV Infection in Bees. The absolute expression levels of 20 genes from Toll (A-D), Imd (E, F) RNAi and two heat shock proteins (G-J), melanization (K-O), as well as Jak-STAT (P-T) pathway after IAPV infection at days 1, 2 and 3 were measured by qPCR. Asterisks indicate significant differences relative to controls (\**P* < 0.05, \*\**P* < 0.01).

**Fig 3.**
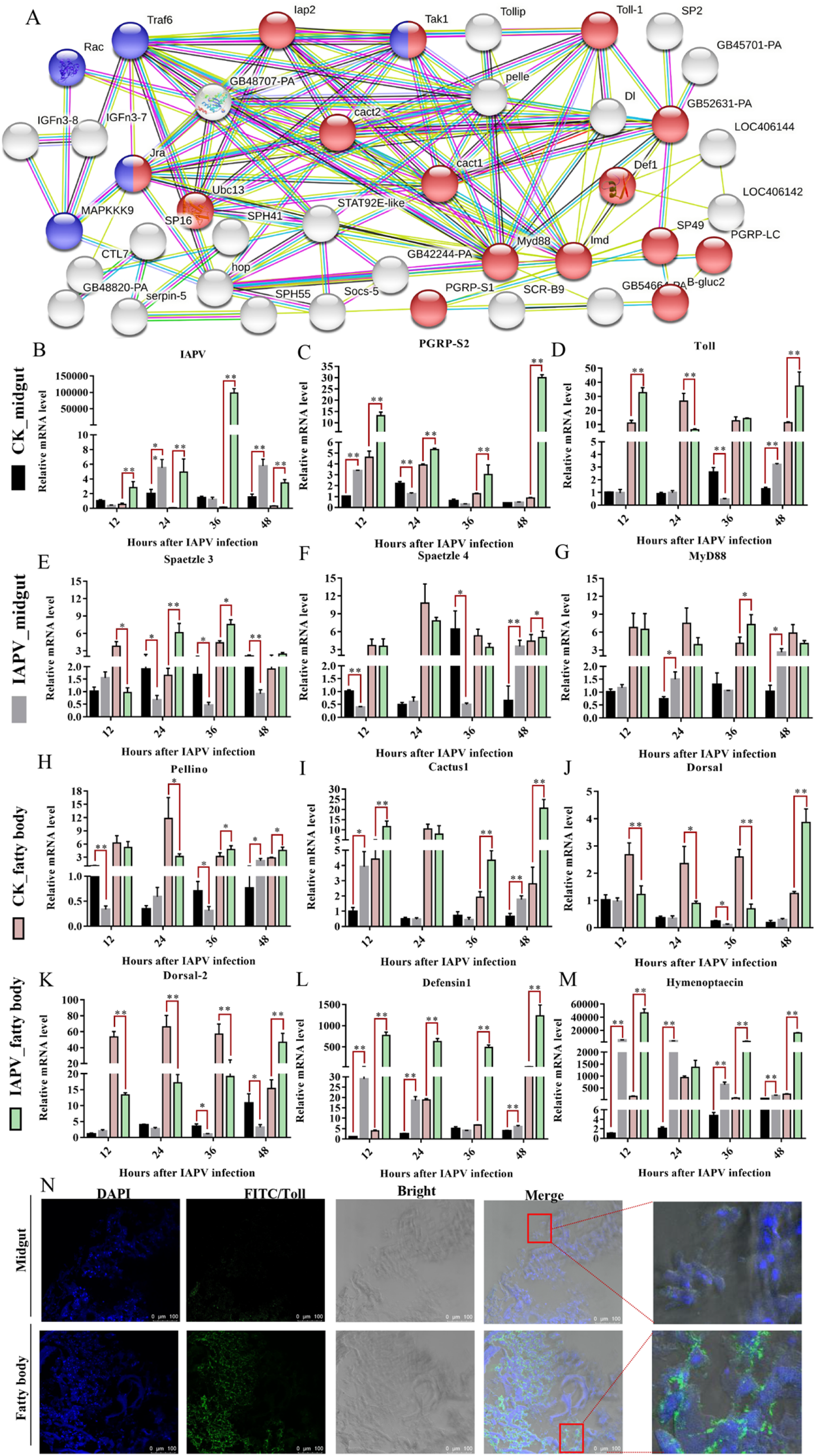
The Toll Pathway is Mainly Activated in the Fat Body. (A) The interaction network overlap of immune genes from Galbrath et al. (2015) and Li-Byarlay et al. (2020) was predicted, and the two major pathways were enriched (*P <0.01*). Red, Toll and Imd signalling pathway (*P<0.00001*); Blue, MAPK signalling pathway (*P <0.01*). The expression of IAPV and main immune genes of Toll pathway in the midgut and fat body at 12, 24, 36 and 48 h post-infection (A-L). (M) Localization of Toll in fat body and midgut using anti-Toll to detect the Toll (in green) and DAPI (blue) at 48 hours post-infection. Asterisks indicate significant differences relative to controls (\**P* < 0.05, \*\**P* < 0.01).

Generally, the midgut, hemolymph and fat body, as the most important immune organ in insects, were also used as a candidate tissue to test the expression of genes confirmed by the results above. As shown in **Fig 3B**, the fat body was more susceptible to IAPV infection than the midgut in *A. mellifera*, suggesting that these AMPs should show higher expression levels in the fat body than in the midgut because they depend on the fat body for transcription. To further confirm the components of Toll involved in the antiviral response, we assessed the expression of all genes known in the predicted Toll pathway, including *Toll, späetzle, MyD88, pellino*, *cactus* and *dorsal* by qPCR (**Fig 3A**). As expected, the *PGRP-S2*, *Toll*, *cactus1*, *hymenoptaecin* and *defensin1* were significantly altered after IAPV infection, and they exhibited a higher expression level in the fat body than that in the midgut (**Fig 3C–M**). Moreover, *späetzle 3, späetzle 4*, *Myd88*, *pellino, dorsal* and *dorsal-2*, to a certain extent, were significantly up-regulated after IAPV infection both in fat body and midgut, especially *späetzle 3* in fat body (**Fig 3C–M**). However, all of the rest genes, such as *cactus 3*, *späetzle 5*, *dorsal-1a* and *dorsal-1b* did not increase (**S2 Fig**). Then, immunofluorescence staining using an anti-Toll polyclonal antibody showed that Toll was localized in the fat body cell membrane but not midgut (**Fig 3N**). These findings suggested that the Toll signalling pathway was the the major response to IAPV infection in the fat body of honey bee.

### Toll pathway activation by PGN drives the production of AMPs against IAPV infection

Due to the lack of honey bee virus cell, we employed the peptidoglycan (PGN from *Staphylococcus aureus*) as a trigger, which had been proved it could activate the Toll pathway and induce the AMPs expression [25]. Since the production of AMPs depends on activation of the Toll or Imd pathway, AMPs expression has been used as a means to determine whether these pathways are activated during infection. The manufacturer’s instruction of PGN stated that conventional final concentrations ranged from 0.1 to 3 μg/mL, so we selected five concentrations (0.05, 0.2, 1, 10 and 50 μg/mL) to investigate the effect of PGN on survivals of honey bees. Similarly, these five concentrations were also applied to test the synergistic effects of PGN and IAPV. Our results showed that all different concentrations of PGN had no impact on the survival of honey bees except a 50 μg/mL (**Fig 4A**). The PGN exhibited the expected results at the concentration of 10 μg/mL when it was mixed with IAPV, and then this concentration was used in the following study (**Fig 4B**). As expected, we found that the PGN (10 μg/mL) not only help the bee to activate the Toll pathway and elicit expression of *PGRP-S2*, *Toll*, *Defensing 1* and *hymenoptaecin* to against IAPV infection (*p < 0.01*) (**Fig 4C–G**), but also reduced the titer of IAPV (**Fig 4H**). Consistently, compared to the IAPV group, the level of expression of these four genes was up-regulated in bees treated with PGN and IAPV all the time, except for days 7, which may be consistent with less IAPV titer at days 7(**S3 Fig**).

**Fig 4.**
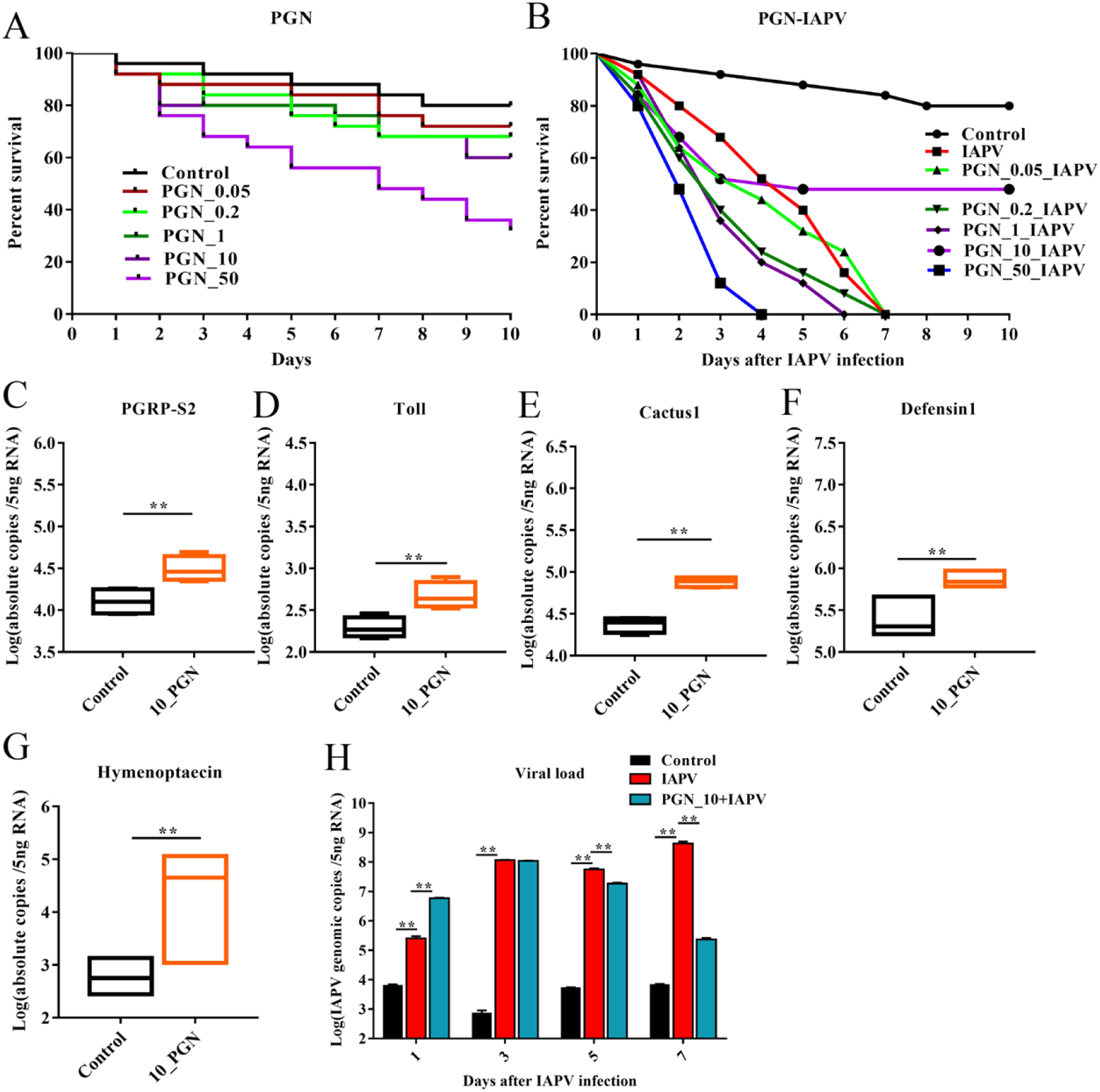
The activation of the Toll pathway by peptidoglycan (PGN, from *Staphylococcus aureus*) Drives the Production of AMPs Against IAPV infection. The effect of PGN at concentrations of 0.05, 0.2, 1, 10 and 50 μg/mL in combination with IAPV on the survival of bees for ten days (A). The effect of PGN at concentrations of 0.05, 0.2, 1, 10 and 50 μg/mL in combination with IAPV on the survival of bees for ten days (B). The activation of Toll pathway 10 μg/mL PGN at days 1, 3, 5 and 7, elicits expression of *PGRP-S2* (C), *Toll* (D), *cactus*1 (E) *defensin*1 (F) and *hymenoptaecin* (G).The activation of the Toll pathway by 10 μg/mL PGN help bees against IAPV infection, the effect of 10 μg/mL PGN on IAPV titer at days 1, 3, 5 and 7 (H). Asterisks indicate significant differences of respective points from the control group (\**P* < 0.05, \*\**P*

### PGRP-S2 is crucial for the Toll pathway activation

*PGRP-S2*, as a member of the PGRP family, showed the most outstanding levels of upregulation, and its expression level was more than 20 times higher than the control at 48 h after IAPV infection in the fat body. Next, to investigate whether PGRP-S2 is crucial for Toll pathway mediated antiviral responses to IAPV infection in honey bee, we silenced *Toll* and each *PGRPs* (PGRP-S1, 2 and 3), respectively, which were significantly changed after IAPV infection, and then measured their expression levels by qPCR. Our results revealed that silencing of *PGRP-S2* (*p* <0.01) and *Toll* (*p* <0.05) significantly reduced the survival of honey bees, and the mass death of bees mainly ranged from days 3 to 5 (**Fig 5A**). Meanwhile, silencing of *PGRP-S2* but not *PGRP-S1* or *PGRP-S3* significantly down-regulated the expression level of Toll (*p* <0.05) (**Fig 5B**). As expected, qPCR and western blot assay results showed, silencing of *Toll* and *PGRP-S2* showed increased susceptibility to IAPV (*p* <0.01) (**Fig 5C–D**) and progressed to inhibit the generation of IAPV structural proteins, VP2 (**Fig 5E**). Notably, the virus abundance was promoted significantly at the protein level in the absence of PGRP-S2 and Toll in a time-dependent manner from days 3 to 5 (*p* <0.01) when IAPV copies were >10^7^ genomes/5ng RNA, similar to the detection limit for IAPV-VP2 (**Fig 5E**). This result was further confirmed by quantitatively assessing the target band intensity (**Fig 5F**). Yet, these inhibition effects were no more until day 6 because the timeliness of dsRNA treatment mainly ranged from days 2 to 6 (**S4 Fig**).

**Fig 5.**
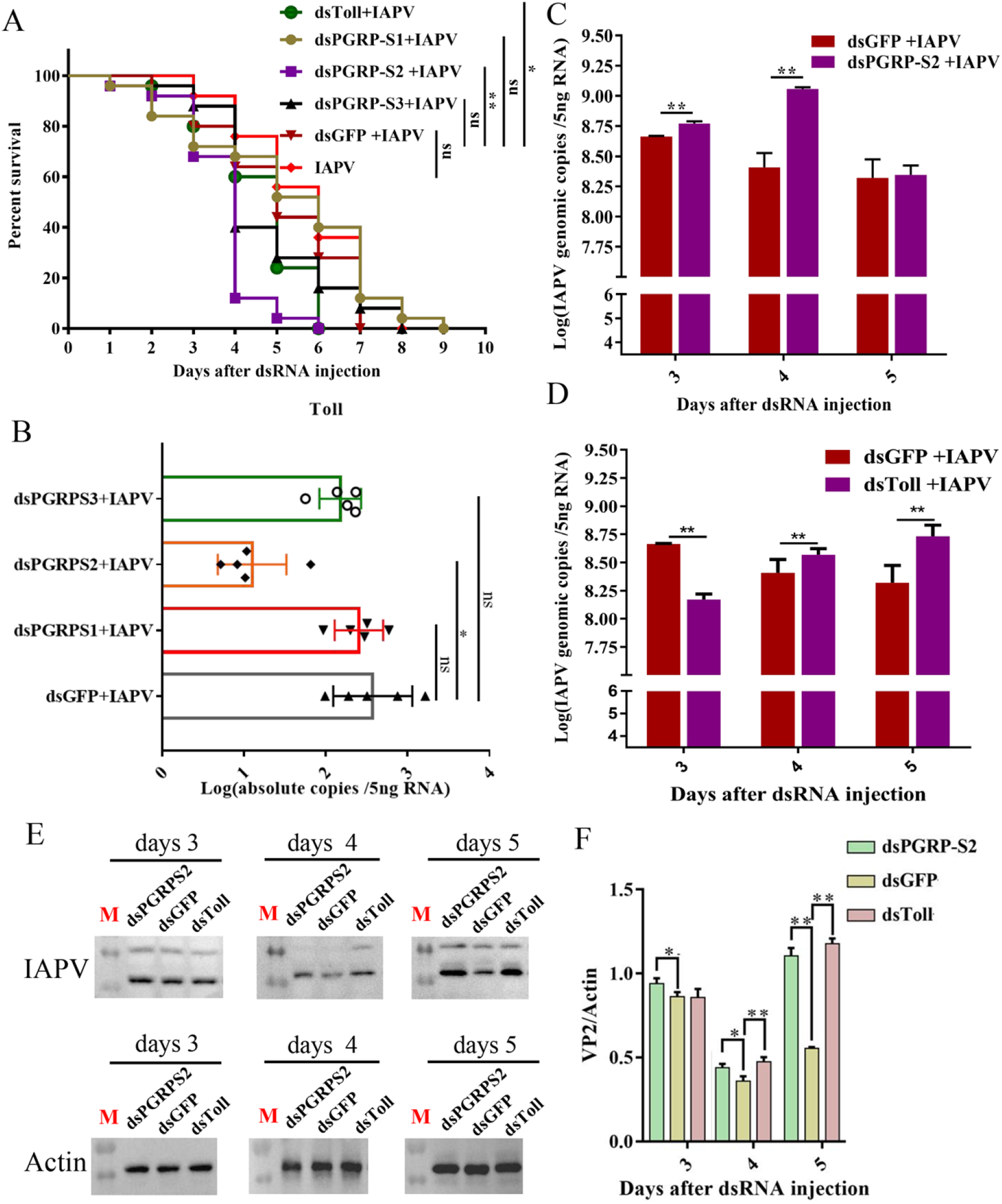
PGRP-S2 is crucial for Toll pathway mediated antiviral responses to IAPV infection in honey bee *Apis mellifera*. (A) The survival rate of bees treated with dsRNA-Toll, dsRNA-PGRP-S1, dsRNA-PGRP-S2 and dsRNA-PGRP-S3. (B) The effect of bees expression level of Toll genes of bees treated with dsRNA-PGRP-S1, dsRNA-PGRP-S2 and dsRNA-PGRP-S3. The effect of IAPV titer over 3 days in bees treated with dsRNA-Toll (C) and dsRNA-PGRP-S2 (D). (E) The effect of IAPV capsid protein abundance over 3 days in bees treated with dsRNA-PGRPS2 and dsRNA-Toll. (F) The relative expression level of IAPV VP2 against actin. Asterisks indicate significant differences relative to controls (\**P* < 0.05, \*\**P* < 0.01).

In addition, on the basis of the above results, we reasoned that PGRP-S2 might induce AMPs expression via the Toll pathway. Therefore, we proposed whether PGRP-S2 and its downstream genes, AMPs, is reduced treatment with dsRNA on *PGRP-S2* and *Toll*. As showed in **S5 Fig**, the dsRNA-PGRP-S2 treated with IAPV-infected bees significantly kept diminishing the induction of *defensin 1* and *hymenoptaecin* expression relative to the bees feeding of unrelated dsGFP with IAPV at days 3, 4 and 5 after dsRNA injection (*P* < 0.01), while the dsRNA-Toll treated with IAPV-infected bees significantly kept reducing the induction of *defensin*1 and *hymenoptaecin* only at days 3 (*P* < 0.01). In addition, the decreased times of *defensin1* both in dsRNA-PGRP-S2 and dsRNA-Toll groups was higher than that of *hymenoptaecin* (**S5 Fig A-D**). What’s more, the dsRNA-PGRP-S1 treated with IAPV-infected bees significantly reduced the expression of *defensin1* only at days 3 (**S5 Fig E**), while the dsRNA-PGRP-S3 treated with IAPV-infected bees significantly facilitated the expression of *defensin1* at days 4 and 5 (**S5 Fig F**). These results suggested an essential function of *PGRP-S2* in IAPV-induced Toll pathway responses in honey bees, and led to the induction of a specific set of genes, including *defensin 1*.

### PGRP-S2 Activated the Toll Pathway via Interacted Directly with VP3 and Toll

The above findings suggested that the Toll signaling pathway was the major response to IAPV infection in honey bee and PGRP-S2 is crucial for the activation of Toll pathway, thus we speculated that co-localization of PGRP-S2 and Toll in the cell surface of the fat body. As shown in Fig **6A–D**, the yellow fluorescence intensity localization on fat body cells was significantly increased in the presence of Toll and PGRP-S2, compared with the mock-infected group. Subsequently, the interaction network between Toll (Toll-1) and PGRP-S2 was predicted by STRING (P <0.0001) (**Fig 6E**). To further verify the interaction between Toll and PGRP-S2, forty-eight hours after IAPV infection, bee samples either mock-infected or infected with IAPV were lysed by RIPA lysis buffer. Then, western blot showed the expression of Toll was up-regulated after IAPV infection, and even IAPV infection accelerated the Toll protein to form dimer and lead to activate the Toll pathway (**Fig 6F**). Subsequently, the co-immunoprecipitation assay was performed using rabbit anti-Toll polyclonal antibody and mouse anti-PGRP-S2 polyclonal antibody, as demonstrated in **Fig 6G**, the interaction between PGRP-S2 and Toll was observed in bee samples either mock-infected or infected with IAPV. However, IAPV infection significantly promoted the interaction in bees, accelerated the expression of PRGP-S2 protein (dimer). Like human PGRP-S [26], PGRP-S2 was also a natural dimer expressed in bees (**Fig 6G**). In addition, the interaction between PGRP-S2 and Toll was specific since Toll did not bind biotinylated IgG (**Fig 6G**).

**Fig 6.**
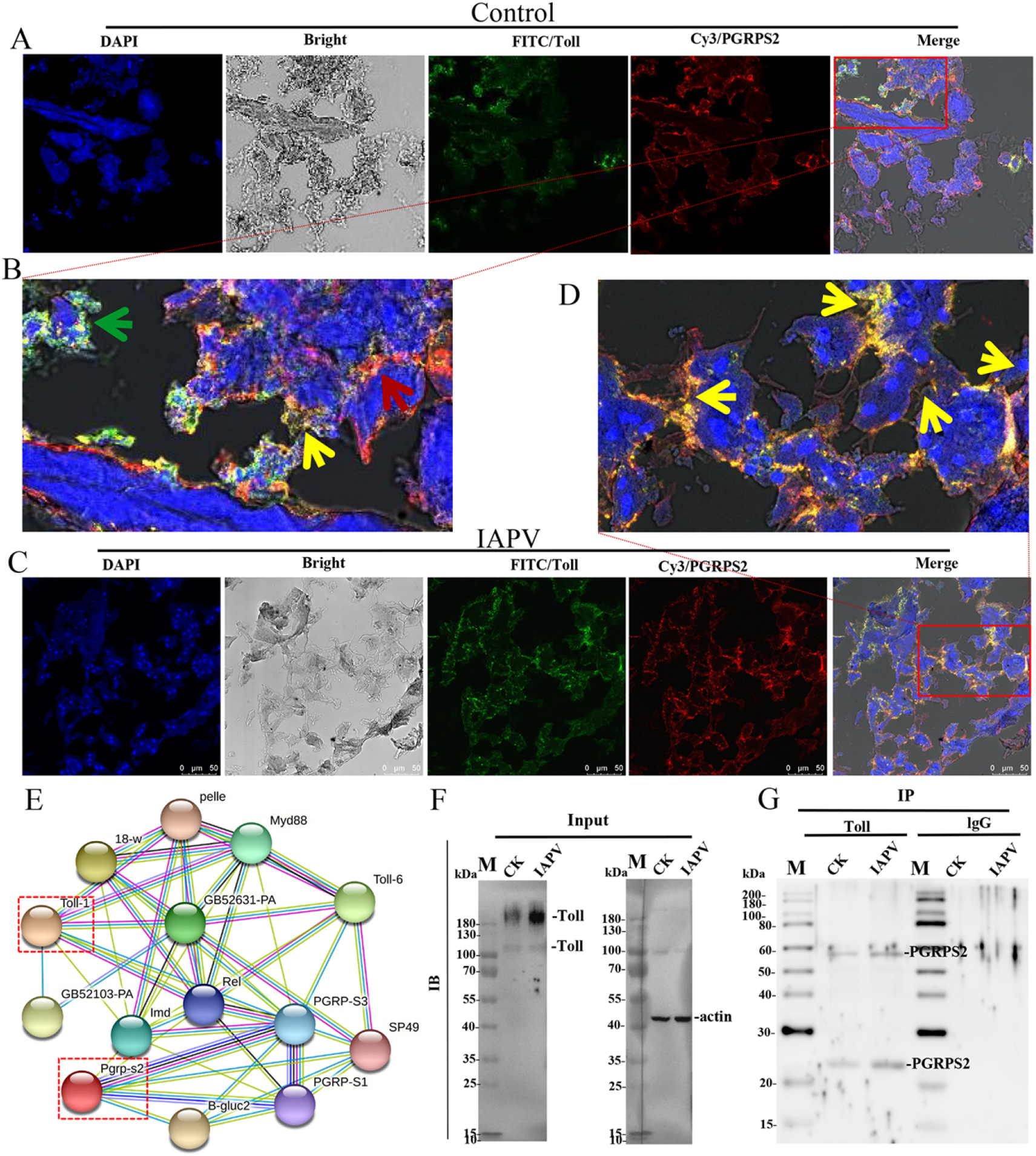
PGRP-S2 Interacts with Toll protein to Mediate Toll Pathway at the Fat body cell member surface. (A) The control 10 bees fat body cells were dissected at 48 hours. Immunofluorescence staining was performed using an anti-Toll monoclonal antibody and mouse anti-PGRP-S2 polyclonal antibody to detect the interaction between Toll (green) and PGRP-S2 (red). Co-localization PGRP-S2 and Toll on the cell surface (yellow). Cells were also counterstained with DAPI to label nuclei (blue). Magnification of the location area is shown in B. Shown is a representative experiment out of at least 100 targeted cells. (C) The 10 IAPV-infected bees fat body cells were dissected 48 hours after infection. Dual staining showed PGRP-S2 and Toll protein colocalization at the cell surface aggregated structures as yellow fluorescence. Cells were also counterstained with DAPI to label nuclei (blue). Magnification of the location area is shown in D. Shown is a representative experiment out of at least 100 targeted cells. (E) The interaction network of Toll and PGRP-S2 was predicted by STRING (P < 0.05). (F) At 48 hours post infection, 10 bees fat bodies (0.01) either mock-infected or infected with IAPV, were lysed by 100 μL RIPA lysis buffer. Total cell lysates were then immunoprecipitated with rabbit anti-Toll polyclonal antibody and mouse anti-actin monoclonal antibody, respectively. (G) About 30 μL supernatant protein complexes (30 μg) from total cell lysates were further precipitated with beads and rabbit anti-Toll polyclonal antibody for 12 hours at 8°C. Protein interaction was detected by immunoblot analysis using mouse anti-PGRP-S2 polyclonal antibodies.

Given that the up-regulated expression of spatzle3/4 and PGRP-S2 was required for activation of Toll pathway after IAPV infection in honey bees, immunofluorescence and Co-immunoprecipitation results also confirmed that Toll pathway was initiated by PGRP-S2 interacting with Toll in honey bee, we sought to determine the possible interaction between PGRP-S2 and capsid proteins of IAPV. According to the genome structure of IAPV (**Fig 7A**), we obtained the soluble PGRP-S2, VP1, VP2 and VP3 proteins of IAPV with His-tagged *in vitro* (**Fig 7B–C**) and identified their specificity using rabbit anti-VP1, -VP2 and -VP3 polyclonal antibody (**Fig 7D**). Co-IP assays and western blot analysis using mouse anti-PGRP-S2 polyclonal antibody with total cell lysates from IAPV-infected bee samples showed that PGRP-S2 can interact with VP3 whereas others can not (**Fig 7E**). We further confirmed this interaction using purified PGRP-S2 *in vitro*, and the signal was more substantial than that of cell lysates (**Fig 7F**). Furthermore, a specific interactioln occurred between PGRP-S2 and VP3 as indicated by the results that PGRP-S2 did not bind with either VP1 or VP2. These data suggest that PGRP-S2 may directly recognize the IAPV via directly interacting with VP3 to active the Toll pathway against IAPV infection in honey bees.

**Fig 7.**
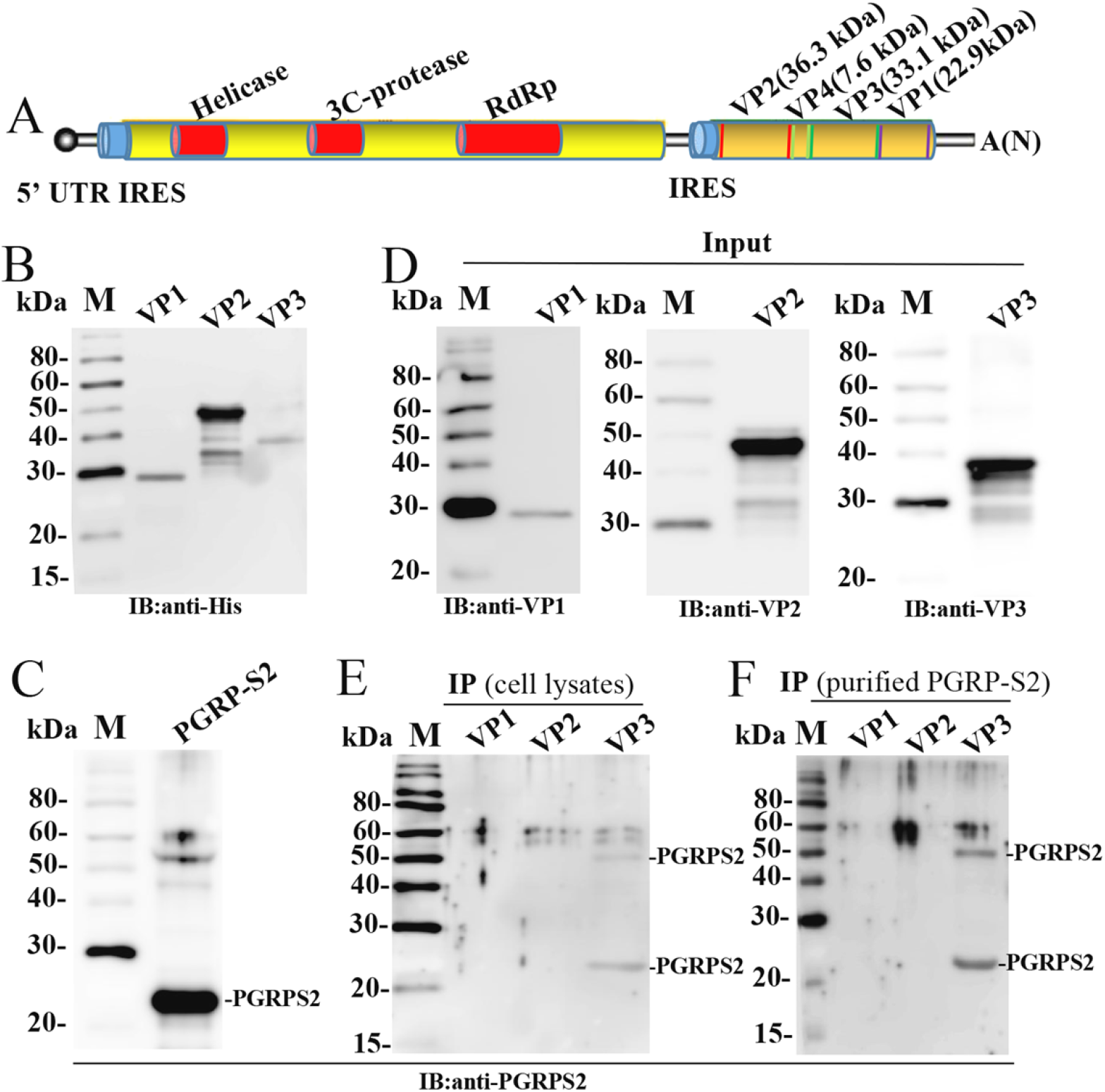
PGRP-S2 Triggers Toll Pathway by Directly Interacting with IAPV VP3 protein. (A) Schematic illustration of IAPV genome structures (RdRp: RNA-dependent RNA polymerase). Total cell was lysed from transformed BL21 strains were further purified with His Pur™ Ni-NTA Resin at 4°C, and then purified IAPV structural proteins were detected by immunoblot analysis using an HRP conjugated anti His-Tag mouse monoclonal antibody (1:1000) (B) and detected for purified PGRP-S2 using mouse anti-PGRP-S2 polyclonal antibody (1:1000) (C). To confirm further the antibody specificity, we used unpurified VP1, VP2, VP3 to detect by immunoblot analysis using rabbit anti-VP1, anti-VP2 and anti-VP3 polyclonal antibody (1:1000) (D). The newly emerging bees was injected with 2 μL IAPV (1012 copies). At 48 hours post-infection, 3 bees (0.1g) infected with IAPV were lysed by 1 mL RIPA lysis buffer. About 30 μL (30 μg) supernatant protein complexes from total cell lysates were further precipitated with beads and rabbit anti-VP1, anti-VP2 and anti-VP3 polyclonal antibody for 12 hours at 8°C. Immunoblot analysis detected bound proteins using mouse anti-PGRP-S2 polyclonal antibody (E). The about 10 μg purified PGRP-S2 was further precipitated with beads and rabbit anti-VP1, anti-VP2 and anti-VP3 polyclonal antibody (about 2 μg) for 12 hours at 8 °C. Bound proteins were detected by immunoblot analysis using mouse anti-PGRP-S2 polyclonal antibody (F).

## Discussion

Although honey bees sense the invading pathogens via signaling pathways to elicit immune response [24], the Toll signalling pathway in response to viral infection in honey bees remains unclear. In this study, we have identified the Toll pathway as a specific-virus immune response of honey bee via a large-scale immune genes screening. Then, we identified and verified PGRP-S2 as a regulator of the Toll pathway in response to IAPV infection. dsRNA-PGRP-S2 and -Toll treated bees exhibited increased IAPV infectivity and mortality, and significantly elevated viral replication after IAPV infection. In addition, PGRP-S2 acted as a PRR through interacting with IAPV-VP3 to induce a PRR effector program that converged on the antiviral Toll pathway. Moreover, these findings provide insights into the way of distinct initiation of the Toll pathway by PGRP-S2 in honey bee.

The essential role for the insect Toll pathway in antimicrobial response is well set up; however, whether the Toll pathway of honey bees serves as antiviral responses have been poorly understood. Honey bee viruses can cause multiple immune responses including Toll pathway but no evidence is associated with activation of this pathway. In this study, we identified and confirmed that the Toll pathway was vital for antiviral response in *A. mellifera* and was in line with that for DXV in *Drosophila* [6], and suggested that Toll pathway is an essential part of the insect antiviral response regardless of the ssRNA and dsRNA viruses. Previous studies demonstrated that Toll pathway was one of the vital regulators of anti-bacterial and -fungi response in *Drosophila,* progressed to regulate the expression of AMPs [27]. However, although the Toll pathway can resist multiple pathogens including bacterial [10], our results showed that the Toll pathway was among humoral immune pathways solely contributing to resist the IAPV infection. Instead, the Toll is activated by the cytokine spätzle, the product of a proteolytic cascade induced upon upstream recognition of bacterial and fungal PAMPs [4, 28–29]. Similarly, our results showed that the process of PGRP-S2 interacting with Toll to activate the Toll pathway to resistance viral infection might require the participation of spätzle 3/4. This conclusion was supported by the fact that *späetzle 3*, *späetzle 4*, *Myd88* and *pellino*, to a certain extent, were significantly up-regulated after IAPV infection both in the fat body as described by the qPCR results as well as increased the viral abundance (**Fig 3**). These results indicated that the Toll pathway might be a distinct antiviral response initiated by specific PGRP. The one critical finding in this study were that honey bee Toll pathway was mainly responsible for fighting IAPV infection. In our study, silencing Toll (Toll-1) exhibited increased IAPV infectivity (**Fig 4**). Although a previous study had shown that Toll-7 could act as an activator of antiviral immunity via autophagy [30], the function of Toll appears to be specific to the antiviral response induced by the Toll pathway was the first evidence for honey bees. The other study showed that the Toll pathway was responsible for resistance to bacterial or fungi in honey bee and virus in *Drosophila* [24, 31]. Yet, our results showed that the Toll pathway was an essential component of viral resistance in honey bee. For example, most of up-and downstream genes of the Toll pathway was significantly regulated than those of other immune pathways after IAPV infection. As shown in **Fig 3A**, transcriptomic analyses from Galbraith et al. have suggested that IAPV infection induced alternation of *hymenoptaecin*, *PGRP-S2*, *PGRP-S1*, *NF-kappa-B inhibitor cactus 1*, *serine protease 14* and *Toll* in honey bee workers after fed IAPV suspension [19]. Also transcriptomic analyses from Li-Byarlay et al. have suggested that the Toll/Imd pathway was significantly enriched in IAPV-infected pupae [20]. However, Chen et al. found that multiple immune biological pathways were significantly up-regulated, especially Jak-STAT pathway [17], which was no different in brood but significantly invoked in adult bees. This indicated that brood and adult bees might use the different immune pathways, or IAPV induced distinct immune responses in adult. For example, a report showed that PGRP-S2 was explicitly induced by adult worker bees but not larvae [32]. Moreover, these discrepancies between Chen et al. and our results prompted us it might be caused by using different IAPVs [17], purified IAPV an infectious clone. Because purified IAPV do not make sure to keep IAPV predominating during the infection, the immune response might be caused by other viruses such as black queen cell virus or sacbrood virus; even IAPV was the primary virus at the start of infection [23]. In addition, although IAPV was similar to acute bee paralysis virus (ABPV) and kashmir bee virus (KBV) in the genome sequence, ABPV infection can not induce humoral and cellular immune responses [33]. These results demonstrated pure virus is essential to explore the immune response of host in the absence of honey bee cell culture system.

The other critical findings found in this study were that PGRP-S2 directly recognized IAPV-VP3, and passed the signal to Toll with the participation of spätzle 3/4 to activate the Toll pathway. Although honey bees have four PGRP genes, three for short and one for long, their differences in expression level between IAPV and control groups prompted us to investigate the role of PGRP-S2 in the honey bee immune response (**Fig 1, 2 and 3**). Silencing PGRP-S2 exhibited increased IAPV infectivity, and honey bee treated with dsRNA-PGRP-S2 displayed significantly elevated viral titer and mortality after IAPV infection compared to the IAPV infection group. Moreover, PGRP-S2 acted as a recognition protein by interacting directly with Toll at the plasma membrane of fat body to induce the Toll pathway, and progressed to resistance to viral infection. Our co-immunoprecipitate assay between PGRP-S2 and Toll displayed an enhanced PGRP-S2-mediated activation of the Toll pathway in a PGRP-S2 dependent manner. Generally, pathogens promoted the dimerization of the Toll/Toll-like receptor, bringing together the corresponding cytoplasmic Toll/interleukin-1 receptor (TIR) domains which also dimerize [34]; our result also showed the IAPV infection accelerated the Toll to dimerize (**Fig 6**). As shown in **Fig 6**, PGRP-S2 was significantly localized in cell surface with Toll and indicated that PGRP-S2 could be a membrane protein. Likewise, PGRP-SD was an extracellular receptor function activating the Imd pathway [35]. Recently, Hou et al. found that honey bee did not have PGRP-SA and Gram-negative binding protein (GNBP) [36]. Previous studies indicated that PGRP-S2 could induce the immune pathways to resistance the fungi and bacterial, while our results demonstrated that PGRP-S2, might be a secreted protein, links viral recognition to Toll pathway and antiviral defence (**Fig 7**). Intriguingly, the interaction between PGRP-S2 and VP3 suggested that IAPV interacted directly with viral structural protein, unlike other viruses that owned glycoproteins such as Vesicular stomatitis virus [37], yet further study need to be confirmed. Previous studies confirmed that PGRP-LC and PGRP-LE could activate the Imd signalling pathway after bacterial infection [38]. PGRP-SD has a vital role in Imd pathway activation for DAP-type bacterial [35]. Contrary to a previous study on the PGRP-LC and PGRP-S2 of honey bee, our data revealed that only PGRP-S2 was constitutive up-regulated after IAPV infection, suggesting that PGRP-S2 has essential yet distinct functions in honey bee immune response to IAPV infection. Together with previous results, *Nosema ceranae* can also induce the up-regulation of *PGRP-S2* in bees [39], and *PGRP-S2* also exhibited increased expression in SBV and DWV infected bees [40], which showed that PGRP-S2 might have a multifaceted function in honey bee immune defence against pathogens.

In addition, quantification analysis on the Toll in midgut and fat body showed that *Toll* gene was mainly enriched in the fat body and had higher expression than in midgut. This was in line with localization results in **Fig 3** and suggesting that Toll was mainly activated in the fat body and then induced the Toll pathway and expressed AMPs. Likewise, the expression of *PGRP-S2* in the fat body was significantly higher than midgut (*p* < 0.01), which was consistent with that of *Bombus ignites* but differed from that of *Bombyx mori* [41, 42]. Contrary to a previous study using the bacterial infection in *Drosophila*, PGRP-LE induced the mainly immune response in the gut [36], our findings did not reveal such role for PGRP-S2. In addition, IAPV infection did not significantly enhance the expression of *PGRP-LC* (**Fig 2**), which is a kind of PRRs similar to *PGRP-LE*. These data indicated that insect might own different recognition receptors in the midgut and the fat body and then trigger the distinct immune pathways responding to different pathogens. However, further studies should focus on whether other *Toll* genes activated and induced the corresponding immune responses in honey bees. What’s more, significant change in *cactus1* prompted us to re-investigate the role of cactus in honey bee immune response to viral infection. In most cases, Dif/Dorsal was considered a transcriptional factor, while our quantification analysis showed that *cactus* had higher level expression than Dif/Dorsal after IAPV infection. Thus, whether cactus had a similar function need to be further explored.

AMPs are the essential effectors of the honey bee immune response to combat viral infection. Both Imd and Toll pathways result in the production of a large number of AMPs, such as *hymenoptaecin* and *defensin* 1. In *Drosophila*, *Dif/Dorsal* were downstream genes of Toll pathway to regulate the *Drosomycin* expression; while our results showed *Dif/Dorsal* mainly modulated the expression of *defensin 1* and *hymenoptaecin*. In line with Wang et al. described that IAPV infection could induce a higher *hymenoptaecin* and *abaecin* expression than other AMPs [43]. Additionally, transcriptomic and epigenomic analysis showed that although about 10 genes involved in Jak-STAT pathway were significantly upregulated in samples between 20 and 48 h, *defensin 1* was up-regulated remarkably at each time point [20]. The increase in *hymenoptaecin* and *defensin1* expression does show that the Toll pathway was caused by IAPV infection. Our RNAi experiments showed that dsRNA-PGRP-S2 and -Toll treatment would increase the bees vulnerability to IAPV infection and then reduce the expression corresponding to downstream genes, such as *defensin 1*. On the contrary, experimental evidence confirmed that *defensin 1* would not be induced by American foulbrood, a pathogenic bacterial of honey bee larvae [44]. These suggested that *hymenoptaecin* and *defensin1* might be specifically induced for IAPV infection. However, this does not mean hymenoptaecin and defensin 1 interacted directly with IAPV capsid proteins due to the IAPV lack the cellular membrane structure [6], and indicated that hymenoptaecin and defensin 1 play a vital role in antiviral response in an indirect manner.

Altogether, the evidence of an antiviral response of the Toll pathway in honey bees obtained in this study provides a typical example of how the innate immune pathway is capable resisting the honey bee virus besides RNAi pathway. The influence of various pathogens on bees immune system should be investigated to understand their mutual relationships, and then would facilitate to development of pharmacological or genetherapy strategies. The activation of the Toll pathway is complex and involves multiple regulatory steps, which are still incompletely understood. Moreover, honey bee has 5 Toll genes and whether other Toll genes are involved in the immune responses of honey bee is unclear. As described by Toll and Toll-7 can induce different immune responses in adult males and females during the vesicular stomatitis virus infection [45]. Our results initially confirmed the existence of fundamental differences in the control of expression of the recognition protein between honey bee and *Drosophila* after virus infection, and then to increase our understanding of the interaction between virus and honey bee, and have implications on honey bee immune responses in functions diversity due to the litter immune genes compared to *Drosophila*.

## Materials and methods

### Honey Bee Samples

All bee colonies were raised using standard beekeeping practices and no typical symptoms of pathogens were observed in these colonies. Experimental bee samples were kept in an incubator at 33°C with 70% relative humidity. All bee samples genotypes used in this study belonged to *Apis mellifera* Ligustica. Honey bee samples were collected from colonies of experimental apiaries of our Institute. A single brood frame with emerging worker bees (*A. mellifera*) was selected from experimental colonies of Institute of Apicultural Research (IAR) and kept them in an incubator at 34 °C with an 80 % relative humidity overnight. Newly emerged bees were obtained after 24 h and used to detect the presence of eight common viruses: Israeli acute paralysis virus (IAPV), Sacbrood virus (SBV), Deformed wing virus (DWV), Black queen cell virus (BQCV), Chronic bee paralysis virus (CBPV), Acute bee paralysis virus (ABPV), *Varroa destructor* virus (DWV-B) and Chinese sacbrood virus (CSBV) (**S2 Table**). The bee samples uninfected with the common viruses mentioned above were placed in groups of 30 bees per small cage for further experiments.

### Generation of an infectious clone of IAPV

Adult IAPV-positive worker bees showing visible paralysis symptoms were collected from IAR experimental apiaries. To purify the virus, IAPV was screened for predominance in the samples, and the absence of other common viruses was also verified. Approximately 50 adult honey bees were collected from 5 colonies of 3 apiaries in which paralysis symptoms were found. Positive samples were used to extract crude virus, which was used to infect healthy honey bee pupae. After three days, IAPV was purified as described by de Mirada [46]. Briefly, an IAPV crude suspension was injected into the abdomen of white-eyed worker honey bee pupae (*A. mellifera*). The injected pupae were raised in an incubator at 34 °C for 3 days. Approximately 260-300 bee pupae were crushed in 40 mL of 0.01 M PBS buffer with 2 mL diethyldithiocarbamic acid. The mixture was centrifuged at 4,000 rpm for 30 min and the supernatant was centrifuged at 24,000 rpm for 4 h. The precipitated sample was resuspended in 1 mL Brjo buffer with a sucrose gradient and centrifuged at 20,000 rpm for 4 h. The virus fraction was recovered in a CsCl gradient and centrifuged at 27,000 rpm for 24 h, and then concentrated the target virus to centrifuge tube using PBS, pH 7.5.

The stepwise cloning strategy for developing the full-length cDNA clone of IAPV was conducted according to Yang et al. [21]. Briefly, viral RNAs were extracted from purified IAPV above using viral RNA kits (QIAGEN) and three fragments covering the full-length genome were synthesized with high-fidelity Phusion HiFi PCR Master Mix (NEB,) using specific primers as shown by Yang et al. [21]. Plasmid pACYC177 (New England Biolabs, Ipswich, MA) was used to clone three fragments and a T7 promoter sequence (TAATACGACTCACTATAGGG) was linked at the 5′ end of the first fragment. The reactions were carried out following the manufacturer’s instructions. All DNA fragments were purified using a QIA Quick gel extraction kit (Qiagen, Germany). Then, the plasmid containing the full-length IAPV was sequenced to verify the sequence stability. As previously described, a standard cloning procedure made IAPV infectious clones [21].

For *in vitro* transcription, the pACYC plasmid containing the full-length cDNA of IAPV was linearized with *Dpn* I, and then, the resulting linearized plasmid was recycled with phenol-chloroform extractions. The HiScribe T7 Quick High Yield RNA Synthesis Kit (NEB) was used to transcribe RNAs *in vitro* following the manufacturer’s recommended protocols and the resultant RNAs were purified with an EasyPure RNA Kit (TransGen Biotech, Beijing). The RNA quality was assessed in a 1.2% gel using formaldehyde denaturing electrophoresis and dissolved in 30 μL RNase-free water. The RNA concentration was determined with a NanoDrop spectrophotometer. For infection, about 2 μL (1×10^12^ genome copies) synthetic RNA of IAPV as described above was injected into a healthy honey bee abdomen with a Hamilton syringe (702) (Hamilton, Switzerland) [21].

In brief, the newly emerging bees were selected and divided into six groups after identifing the negative for IAPV and other common viruses mentioned above; each small cage contained 30 individuals placed in an incubator as previously described [47]. Treated bees were infected with 2 μL of purified synthetic IAPV RNA (about 1×10^12^ genome copies) (**S1C Fig**), and the control group was injected with phosphate buffer (PBS). After 24 h, the IAPV group was treated with sucrose (50%) until the end of the experiment, and then all groups were transferred to an incubator at 30 °C and 60% relative humidity. Dead bees were removed and transferred to an Eppendorf tube and they were immediately frozen at −80 °C until analyses. Each group consisted of three replicates. To detect the IAPV replication, did the tagged qRTPCR perform

### Selection of immune genes

To identify possible immune pathways in regulating related genes expression in response to IAPV infection, we firstly selected 16 immune genes according to the descriptions of Evans et al. [24] and Chen et al. [17]. Then, to further characterize the immune pathways responsible for resisting the viral infection, we selected 20 immune-related genes involved in different immune pathways, including the Toll pathway, Imd pathway, Jak-STAT pathway, and RNAi and melanization pathway as shown in **S1**, **S3 and S4 Table**.

### qPCR

Total RNA was extracted with the TRIzol Kit (Ambion, Life Technologies, USA) following the manufacturer’s instructions. A NanoDrop 2000 (Thermo Scientific, United States) was used to check the quality of each RNA sample. The RNAs quality with their A260/280 ratios ranging from 2 to 2.2 were used to synthesize cDNA using the PrimeScript RT Reagent Kit with gDNA Eraser and mix primers according to the manufacturer’s instructions. Next, PCR was performed using specific forward and reverse primers according to the corresponding to the primers’ action conditions (**S3 and S5 Table**). The cDNAs of all samples were stored at −20°C until use.

Individual bee was cut into two parts in dorsal midline, and half of the bee was used for qPCR and the other half for western blot. cDNA produced by conventional RT-PCR was subjected to real-time quantitative PCR using a LineGene9600 instrument. The primers used for detecting the IAPV replication was as follows: IAPV Forward primer of 5′-:CCAGCCGTGAAACATGTTCTTACC-3′ and Reverse primer of 5′-ACATAGTTGCACGCCAATACGAGAAC-3′ were used to amplify a 167-bp fragment; housekeeping gene forward and reverse primers of β-actin (5′-TTGTATGCCAACACTGTCCTT-3′ and 5′-TGGCGCGATGATCTTAATTT-3′) were used to amplify the reference gene [48]. To assess the possible immune genes response to IAPV infection, we analyzed the transcript level of 16 immune genes from different immune pathways according to the results of Evans et al. and Chen et al. [24, 17], and then further identify the immune pathways on another 29 genes using the specific primer pairs shown in **S3 Table**. Standard curves for absolute quantification of IAPV load and immune genes were shown in **S4 Table.** qPCR was performed using the KAPA SYBR FASTqPCR kit Master Mix (Sigma-Aldrich, USA) according to the manufacturer’s instructions. The cycling protocol was 95 °C for 3 min followed by 40 cycles of 95 °C for 3 s, 60 °C for 30 s, and 72 °C for 20 s. Standard curves were prepared by performing real-time qPCR with serial 10-fold dilutions of known concentrations of the gene-specific amplicons, approximately 100-300 bp in length. The melting curve dissociation was analyzed to confirm each amplicon. The concentration of nonspecific primer amplification was measured by performing the RT reaction in the absence of added primers followed by specific-primer quantitative real-time PCR and was found to be negligible under our working conditions. The obtained results were analyzed using the 9600 Plus Software.

### Antibodies preparation

Mouse monoclonal antibody to β-actin was purchased from Abbkine (USA). HRP conjugated anti His-Tag mouse monoclonal antibody was prepared by Cwbiotech. The rabbit polyclonal antibodies against a peptide corresponding to IAPV structural protein VP1, VP2 and VP3 were synthesized by Hangzhou HuaanBiotech. The rabbit polyclonal antibodies against the peptide corresponding to 807-1068 aa of the Toll (XM_016911914.1) was synthesized by ABclonal Technology and got the 2 mg/mL purified antigen through Ni-IDA (GE Healthcare, USA). The mouse polyclonal antibodies against two parts of PGRP-S2 (NM_001163716.1), GDSLYELIKTWPHWSSI-C and RALAQNPPPFVIIHHSATDSC were synthesized respectively by ABclonal Technology, and these two polypeptides were linked with KLH, and then to immunized mice.

### Western blotting

Total proteins were extracted from halves of bees as mentioned above by RIPA lysis buffer according to the manufacturer’s instructions. The bees were ground with steel beads in PBS containing 0.2% DTT and 1 μM PMSF. An additional 200 μL extraction buffer was added to the resulting protein extracts and quickly centrifuged. Then, the supernatant was centrifuged for 10 min at 12,500 rpm at 15 °C, and 200 μL of chloroform was added and centrifuged again at 4 °C at 12,500 rpm for 10 min. The protein was resuspended in DDW and diluted with 2× concentrated sample buffer (100 mM Tris–HCl, pH 6.8, 4% SDS, 17% glycerol, and 0.8 M 2-mercaptoethanol), boiled for 3 min and subjected to sodium dodecyl sulfate-polyacrylamide gel electrophoresis (SDS-PAGE) at a constant voltage (110 V). The polypeptides were separated on 12% SDS-PAGE before electrophoretic transfer to nitrocellulose membranes (Pall Corporation, New York, USA). For colloidal Coomassie staining, the gels were fixed for 30 min in 0.85% o-phosphoric acid/20% methanol and then stained in 20% methanol overnight according to the manufacturer’s instructions. The gels were destained in 25% methanol. To detect IAPV antigens, the blots were blocked with 5% skim milk powder and probed with different rabbit polyclonal antibodies against IAPV three structural proteins synthesized by Hangzhou HuaanBiotech. The blots were developed using the Amersham ECL plus reagent (GE Healthcare, Chalfont St Giles, GB) and exposed to a C-Digit scanner (Licor, Lincoln, Nebraska) and imaged by photography. Similar methods were employed to detect the antigens of PGRP-S2 and Toll.

### RNA silencing

dsRNAs were generated and used essentially as previously described by Deng et al. [48]. Briefly, preparation of DNA templates of 7 immune-related genes and GFP were performed with gene-specific amplicons, approximately 300-500 bp in length. Gene-specific primers for 7 immune-related genes (*Toll, PGRP-S1, PGRP-S2, PGRP-S3, cactus 1, cactus 2 and cactus 3*) and GFP were designed to include T7 promoter sequences (TAATACGACTCACTATAGGGAGA) at the 5′ ends of each strand and were subsequently used to amplify target fragments using PCR from cDNA (S4 **Table**). The PCR product was used to the transcript in vitro using MEGAscript RNAi Kit (Invitrogen, USA) to generate RNA strands at 37°C for 4 h. Subsequently, the reaction product was subjected to DNase and RNAse digestions, incubated at 37°C for 1 h, and then purified them with RNAeasy and quantified with a spectrophotometer (NanoDrop Technologies, Wilmington, DE, United States). Then, 30 newly emerging bees in every group were confirmed to be negative for common viruses, and then for further experiment and three repetitions for each treatment. Each of dsRNA (5μg) was injected into newly emerged worker bees of *A. mellifera* at 0 and 48 h, respectively. Honey bees were infected with 2 μL IAPV (~10^12^ genome copies) at 24 h after dsRNA bathing and assayed at the indicated time point post-infection. The interference efficiency was determined by qPCR at 24-, 48-, 72-, 96-, 120- and 144-hours post-injection. dsGFP was used as the negative control.

### Inducement of the Toll pathway by peptidoglycan (PGN)

β-glucans and lysine-type peptidoglycan are thought to be the primary triggers for the Toll pathway of fungi and most Gram-positive bacteria [49, 50]. In addition, it has been reported that PGN of *Staphylococcus aureus* can induce the Toll signalling pathway via enhancing RANKL-stimulated proteins expressions of Toll-like receptor 2 [51]. Therefore, we used PGN as a trigger to confirm the responses of the Toll pathway, and then to investigate the role of this pathway in anti-IAPV. The newly emerged worker bees were divided into 12 groups **to end this**. Each of them contained 30 individuals and was fed with a 50 % (W/V) sucrose solution mixed with PGN (Sigma, USA) at different concentrations (0, 0.05, 0.2, 1, 10 and 50 μg/mL) for 10 days. Then, different concentrations of PGN (0, 0.05, 0.2, 1, 10 and 50 μg/mL) were selected to investigate its role against IAPV. The IAPV-treated groups were injected with purified synthetic viral RNA; and the control groups were fed with 50 % sucrose every day. After 24 h, the IAPV groups were treated with sucrose (50%) containing 6 different concentrations of PGN (0, 0.05, 0.2, 1, 10 and 50 μg/mL) until the end of the experiments. Each concentration group, as well as the control group, consisted of three replicate. Dead bees were removed and transferred to an Eppendorf tube and were immediately frozen at −80 °C until analyses. All above experiments were run in three replicates.

### Tissue dissection and scanning electron microscopy examination

To verify the immune response in different immune tissues, five bees were collected from each of three groups maintained in incubator at 48 hours post infection with IAPV. Under a dissecting microscope, bee samples were fixed on the wax top of a dissecting dish with insect pins, and then were cut along the dorsal midline with scissors from the tip of the abdomen to the head. The scissors and forceps were washed with 0.3% sodium hypochlorite three times, followed by a final rinse in sterile water after isolating each tissue. The fat body and midgut were carefully removed from each bee. Then, the dissected tissues were washed in 1×PBS buffer (10 mM, pH 7.4). Finally, as shown in S**6 Fig,** fat body and midgut were observed according to the method reported by Letzkus et al. With scanning electron microscopy (SEM, Hitachi S-4800 SEM system, Hitachi High-Technologies Corporation, Japan)[52].

### Immunofluorescence (IF)

Fat body and midgut of 5 honey bees were dissected and removed as above, and then were fixed with 4% formaldehyde for 60 min as previously described methods (Chen et al., 2014), followed by frozen with tissue embedding agents (sakura, Japan) at −20°C. Subsequently, 5μm frozen sections of honey bee fatty body and midgut were cut and affixed onto glass slides designed to enhance tissue adhesion (Platinum pro; Matsunami Glass Ind., Kishiwada, Japan). Slides were sequentially washed for 10 min with distilled water and PBS (pH 7). To keep the autofluorescence intensity, we recycled antigens to enhance the intensity of the primary antibody binding. Briefly, we boiled slides with target cells for 3 min in 0.1 M citrate buffer and then were allowed to spontaneously cool to room temperatures under the same buffer. Next, the sections were soaked in the washing buffer (0.1% Tween 20 in PBS) for 15 min and slides were blocked for 1 h with PBS buffer containing 5% milk at room temperature. The primary antibodies (anti-Toll antibody from rabbit, an anti-PGRPS2 antibody from mouse; dilution: 1:200) was diluted in a block and incubated with the sections overnight at 4 °C and with fluorescein isothiocyanate (FITC)-conjugated goat anti-rabbit IgG and Cy3-conjugated goat anti-mouse IgG (Proteintech Group) for 1 h at the room temperature. Then, the cells were washed three times with PBST and incubated in the secondary antibodies:anti-rabbit antibodies (AlexaFluor488, Invitrogen, USA) (dilution, 1:200) and goat anti-mouse lgG (H+L) Cy3 conjugate antibody (Proteintech, USA) (dilution: 1:100), for 2 h at room temperature. After being rinsed three times in 1xPBS, nuclei were stained with 4’,6-diamidino-2-phenylindole (DAPI) (Roche) for 4 min at room temperature and observed them under a Laserscaing Confocal Microscopy (TCS SP8 STED, Leica, German). The fat body and midgut stained cells were analyzed with automated image analysis software by a confocal laser scanning microscopy (FluoView FV1000; Olympus, Tokyo, Japan). Experiments were performed three times.

### Expression and purification of IAPV capsid proteins and PGRP-S2

To obtain soluble PGRP-S2 and 3 structural IAPV proteins of VP1, VP2 and VP3 *in vitro*, plasmids of pET28a-PGRP-S2, pET28a-VP1, pET28a-VP2, pET28a-VP3 were constructed by homologous recombination with a ClonExpressMultiS One-Step Cloning Kit (Vazyme Biotech Co.) according to the manufacturer’s manuals. In brief, total RNA were isolated from 3 IAPV-infected bees and followed by cDNA syntheses with a reverse transcription kit. These cDNAs were used to amplify full-length of PGRP-S2, VP1, VP2 and VP3 with high-fidelity Phusion HiFi PCR Master Mix (NEB, Ipswich, MA, USA) using specific primers (**S6 Table**). PCR products were inserted into pET28a by a homologous recombination; and transformed into *E. coli* BL21 strain (CWBIO). The correct recombinant positive single colony of *E. coli* BL21 shuffling pET28a-PGRP-S2, pET28a-VP1, pET28a-VP2, or pET28a-VP3 were added to LB medium (10 mL) with 50 μg/mL kanamycin and incubated overnight. Then, 250 mL of LB was inoculated with 2 mL of the overnight culture and incubated until the OD 600nm reached 0.5 at 37 °C. After that, protein expression was induced by 0.3 mM isopropyl-β-D thiogalactopyranoside (IPTG) at 16 °C for 24 h.

For purification of the fusion proteins by His Pur™ Ni-NTA Resin (Thermo Scientific, Beijing, USA), the 250 mL IPTG-induced bacterial solution was centrifuged at 4 °C at 12000 rpm for 10 minutes, and the pellet was resuspended in 25 mL 1X PBS (pH7.4). About 25 mL filtrated supernatant contained targeted soluble protein was added to the Gravity column with 1 mL His Pur™ Ni-NTA Resin, and then the protein was eluted with 5 mL different concentrations imidazole (25, 50 250 and 500 mM). Low concentrations of imidazole washed unbound miscellaneous proteins and the fused targeted protein was collected in 5 mL 250 mM imidazole. Supernatant soluble protein concentration was determined by BCA Protein Assay Kit (CWBIO) according to the instructions. Finally, the targeted protein was separated by SDS-PAGE and identified by Western blot using HRP Conjugated Anti His-Tag Mouse Monoclonal Antibody (1:1000, CWBIO).

### Co-Immunoprecipitation (Co-IP)

To verify the interaction between PGRP-S2 and IAPV capsid proteins, or Toll, we used the Pierce™ Classic Magnetic IP/Co-IP Kit (Invitrogen, USA) as a system for enabling highly effective and efficient antigen Co-IP following the manufacturer’s instructions. Briefly, the 30 newly emerging bees were injected with 2 μL IAPV (10^12^ copies), and then 10 bees fat body (0.1g) either mock-infected or infected with IAPV 48 hours were lysed by ice-cold IP Lysis/Wash Buffer and incubated for 30 minutes on ice through periodic mixing, and the lysate was centrifuged 10 min at 12000rpm 4 °C. Next, 30 μL supernatant lysates (about 30 μg) were immunoprecipitated using 2-5 μg of rabbit anti-Toll polyclonal antibody as a trap in a 1.5 mL microcentrifuge tube and the rabbit IgG was used as the control blank antibody. The mixture was incubated for 1-2 h at room temperature. Subsequently, adding 25μL (0.25 mg) of Pierce Protein A/G Magnetic Beads into action tubes to bind antibodies. Next, the mixtures of antigen and antibody was added to the tubes containing pre-washed magnetic beads and incubated at room temperature for 1 h. A 500 μL of IP Lysis/Wash Buffer was also added to the tubes and gently mixed, and then collected the beads and discarded the supernatant. Then, put 100 μL of Lane Marker Sample Buffer (diluted five-fold with purified water) into the tubes and heated the samples at 96-100 °C for 10 min in the heating block. Finally, the beads magnetically separated and saved the supernatant-containing target antigens were detected with mouse anti-PGRP-S2 polyclonal antibody by Western blot. The interaction between IAPV capsid proteins and PGRP-S2 was verified using a similar method. In brief, about 10 μg purified VP1, VP2, VP3, PGRP-S2 and 2 μg rabbit anti-VP1, anti-VP2 and anti-VP3 polyclonal antibodies were used in the following experiments. The interaction networks of corresponding proteins of targeted genes were predicted using STRING (https://cn.string-db.org/).

### Statistical analyses

The average longevity among the different treatment groups was statistically tested with a Kaplan-Meyer survival analysis with Log Rank Test Statistics using the R statistical software. The variation in the IAPV titer between the treatments was analyzed using one-way ANOVA. The qPCR data were tested for normality with a Kolmogorov-Smirnov test and log-transformed if they did not show normality (**S5 Table**). For fold changes of immunity genes, the values less than one indicate significant transcriptional down-regulation, while those higher than 1 indicate up-regulation. One-way ANOVA was used to analysis the statistical significance of calculated indicators between each treatment and the control, and the mean gene expression was compared by Tukey’s multiple comparisons using GraphPad Prism 7.0. *P* < 0.05 and *P* < 0.01 represent significantly and extremely different at the 0.05 and 0.01 levels. The grayscale analysis of Western blotting bands between dsRNA treatment and the control group was measured using grayscale image analysis in ImageJ platform (version 1.52, MD, USA). The value of grayscale among the different groups was calculated and Tukey’s multiple comparison test analyzed the significant variation.

## Supporting information

**S1 Fig. The proliferation of IAPV**. (A) Survival rates of the newly emerging bees for ten days after IAPV infection. (B) Identified the pathogenesis of IAPV by qPCR. The injection (C) and caused the typical symptoms (D).

**S2 Fig. Anti-IAPV function of Toll pathway mainly activated in fat body.** The expression levels of Toll pathway genes *späetzle 5* (A), *cactus 3* (B), *dorsal-1a* (C) and *dorsal-1b* (D) in midgut and fatty body of bees either mock-infected or infected with IAPV at 12 hours, 24 hours, 36 hours and 48 hours after IAPV infection. Asterisks indicate significant differences relative to control (\**P* < 0.05, \*\**P* < 0.01).

**S3 Fig. The Activation of Toll pathway by peptidoglycan (PGN) Drives the Production of AMPs to Against IAPV infection.** The activation of Toll pathway by IAPV and 10 μg/mL PGN with IAPV at days 1, 3, 5 and 7, elicits expression of *PGRP-S2* (A), *Toll* (B), *defensin1* (C) and *hymenoptaecin* (D). Asterisks indicate significant differences of respective points from control group (\**P* < 0.05, \*\**P* < 0.01).

**S4 Fig. The timeliness of RNAi on target gene expression over 6 days post infection with IAPV.** Silencing effects on *PGRP-S1* (A), *PGRP-S2* (B), *PGRP-S3* (C), *Toll* (D).

**S5 Fig. The effect of bees treated with dsRNA-PGRP-S2 or dsRNA-Toll on the downstream genes.** dsRNA-PGRP-S2 treatment restrains the expression of downstream immune *defensin 1* (A), *hymenoptaecin* (B) at day 2, 3, 4 and 5 after dsRNA injection. dsRNA-Toll treatment restrains the expression of downstream immune *defensin1* (C) and *hymenoptaecin* (D) at days 3, 4 and 5 after dsRNA injection. Asterisks indicate significant differences relative to control (\**P* < 0.05, \*\**P* < 0.01).

**S1 Table. Honey bee central immune genes and pathway against IAPV**

**S2 Table. Primers are used to detect honey bee common viruses.**

**S3 Table. List of primers for qPCR.**

**S4 Table. Standard curves for absolute quantification of IAPV load and immune genes.**

**S5 Table. Primers used to prepare the dsRNA**

**S6 Table. Primers were used for amplification of three structural protein genes and PGRP-S2.**

## Acknowledgments

We would like to express our thanks to Dr. Hong Zhang, Huipeng Yang, Yueqin Guo, Kai Wang and Jiangli Wu for their value assistant in the microscopy. This research was fund by The National Natural Science Foundation of China (31572471, 31811530276) and the Fundamental Research Funds for CAAS (Y2020PT17), the Agricultural Science, Technology Innovation Program of CAAS (CAAS-ASTIP-2020-IAR), GDAS Special Project of Science and Technology Development (2018GDASCX-0107), The Key Technology Research and Development Program of Guangxi province (AB16380094).

## Author Contributions

Conceptualization: Chunsheng Hou.
Data curation: Yanchun Deng, Sa Yang.
Formal analysis: Yanchun Deng, Sa Yang, Hongxia Zhao, Ji Luo, Zhiqiang Lu.
Funding acquisition: Chunsheng Hou.
Investigation: Yanchun Deng, Sa Yang, Hongxia Zhao, Ji Luo.
Methodology: Yanchun Deng, Sa Yang, Hongxia Zhao, Ji Luo, Zhiqiang Lu.
Project administration: Chunsheng Hou.
Resources: Chunsheng Hou.
Software: Yanchun Deng, Sa Yang.
Supervision: Chunsheng Hou.
Validation: Chunsheng Hou, Yanchun Deng, Sa Yang, Zhiqiang Lu.
Visualization: Chunsheng Hou, Yanchun Deng.
Writing – original draft: Chunsheng Hou, Yanchun Deng.
Writing – review & editing: Chunsheng Hou, Yanchun Deng, Sa Yang, Zhiqiang Lu.

## Data Availability Statement

All relevant data are within the manuscript and its Supporting Information files.

